# Effect of nebulised BromAc^®^ on rheology of artificial sputum: relevance to muco-obstructive respiratory diseases including COVID-19

**DOI:** 10.1101/2021.12.28.474344

**Authors:** Krishna Pillai, Ahmed H. Mekkawy, Javed Akhter, Sarah J. Valle, David L. Morris

## Abstract

Respiratory diseases such as cystic fibrosis, COPD, bronchiectasis asthma and COVID-19 are difficult to treat owing to viscous secretions in the airways that evade mucocilliary clearance. Since earlier studies have shown success with BromAc^®^ as mucolytic agent for treating a rare disease known as pseudomyxoma peritonei (PMP), we tested the formulation on two gelatinous airway representative sputa models, in order to determine whether similar efficacy exist.

The sputum (1.5 ml) lodged in an endotracheal tube was treated to aerosolised N-acetylcysteine, bromelain, or their combination (BromAc^®^) using a nebuliser with 6.0 ml of the agents in phosphate buffer saline, over 25 min. Controls received phosphate buffer saline. The dynamic viscosity was measured before and after treatment using a capillary tube method, whilst the sputum flow (ml/sec) was assessed using a 0.5 ml pipette. Finally, the sequestered agents (concentration) in the sputa after treatment were quantified using standard bromelain and N-acetylcysteine chromogenic assays.

Results indicated that bromelain and N-acetylcysteine affected both the dynamic viscosities and pipette flow in the two sputa models, with changes in the former parameter having immense effect on the latter. BromAc^®^ showed a greater rheological effect on both the sputa models compared to individual agents. Further, correlation was found between the rheological effects and the concentration of agents in the sputa.

Hence, this study indicates that BromAc^®^ may be used as a successful mucolytic for clearing airway congestion caused by thick mucinous immobile secretion, however further studies with patient sputum samples using aerosol BromAc^®^ is warranted.

## Introduction

In the healthy state, secretions in the respiratory tract are continuously cleared by the cilia lining the air passages and expectorated as sputum. However, this is both dependent on the constituency and rheology of the secretion which affects ciliary clearance [1, 2]. Viscosity in the range 12-15 Pa and an elastic modulus of 1 Pa are necessary for optimal mucocilliary clearance [3, 4] Since the airways are continuously exposed to dust particles, pathogens and other exogenous molecules, the secretion propelled by ciliary motion serves to expel these harmful molecules continuously [5] and any disruption may result in accumulation leading to respiratory infection, pneumonia etc. [6]. In cystic fibrosis, a genetic disorder of the CFTR (cystic fibrosis transmembrane conductance receptor) gene, thick mucinous secretion accumulates in the respiratory tract resulting in mucus plugging, bacterial infection and progressive deterioration in lung function [7]. A similar accumulation of mucinous secretion with bacterial infection may result in chronic obstructive pulmonary disease (COPD) and other respiratory infectious diseases [8]. In fatal asthma, airway plugging inevitably with mucus-rich sputum is almost seen [9]. Hence, stasis of airways mucus secretion may lead to a variety of respiratory disorders.

Normal airway secretion is composed of mucins (MUC5B and MUC 5AC), water, sodium chloride, bicarbonate, and cellular materials with a pH of 7.0 [10-12]. The mucins serve as a protective barrier for the epithelial cells lining the respiratory tract as well as tissues beneath. In a diseased state, such as bacterial infection, owing to inflammation excess mucin is secreted by the goblet cells as a protective measure with infiltrating white blood cells [13], accumulation of DNA fragments from dead cells and other solids along with sodium, chloride and bicarbonate imbalance [14] that results in loss of water with the secretion becoming thick and purulent (bacterial infection) that eventuates in the reduction or absence of ciliary clearance [15]. Acidic pH in cystic fibrosis sputum as reported in some studies [16], may encourage the cross linking of DNA, mucins and cellular fragments with additional milieu for bacterial growth and sputum stasis [17].

Recent studies have shown a great similarity between the respiratory secretions of COVID-19 patients and cystic fibrosis (CF) indicating that successful treatment to mobilise the sputum in CF may be used for treating COVID-19 [18] Elevated Muc1 and Muc5AC are seen in COVID-19 sputum [19]. Mucus production stimulated by IFN-AhR signalling triggers hypoxia in a mouse model of COVID-19 [20]. Lower airway viscous secretions in COVID-19 lead to the requirement for a high inspiratory plateau pressure, which required suction pressure of 20 KPa causing closure of distal airways [21]. Stickiness of sputum in associated with more severe disease in COVID-19 patients [22]. Cytokine storm and sudden mucus hypersecretion is well understood to be an important part of COVID-19 for which one drug is inadequate [23]. Further elevated levels of solids including proteins, hyaluronic acid, double stranded DNA (dsDNA) and infiltrating cells along with bacterial colonization was observed in both the diseases as compared to normal sputum, without significant changes in pH (7-8) although some studies on CF have shown a depressed pH [15]. Elevated level of solids has also been shown to have a considerable effect on the rheology of the secretion and hence its clearance. Noticeably, the concentration of double stranded (dsDNA) is much higher (about 600 µg/ml) as compared to hyaluronic acid (about 7.0 µg/ml) in COVID-19 lung secretion suggesting that dsDNA may have a higher impact on the rheology of the secretion.

Treatment options to enhance the clearance of stagnating sputa in CF is through rehydration using saline [24], agents such as Acetylcysteine (NAC), L-cystine, cysteamine and other pharmaceutical agents that are mucolytics [25-27] together with additional antimicrobial agents when there is infection [28]. In diseases such as cystic fibrosis, if stasis of the secretion in the respiratory tract can be halted or modulated, then the invasion of microbes may be reduced or avoided [29]. Similarly in diseases such as pneumonia, COPD and CF, the prevention of muco-ciliary stasis may enable the prevention of microbial invasion since stasis leads to accumulation of foreign agents including microbes. Prevention of respiratory secretion stasis in COVID-19 may prevent deterioration in lung function, transplantation, and death. Stasis of thick pulmonary secretions prevents oxygenation with rapid respiratory failure in COVID-19 patients [22, 30]. Further, with current development of various therapeutic agents for nasal delivery or through the respiratory route, clear airway passages are a requirement for such treatment.

BromAc^®^, a combination of Bromelain and Acetylcysteine has shown both anticancer and mucolytic properties [31-33] and is currently undergoing phase 2 clinical evaluation for the treatment of pseudomyxoma peritonei (PMP) where cancer cells secrete copious amounts of mucin in the peritoneal cavity that eventually restricts nutritional intake resulting in death [34]. Since both the agents in BromAc^®^ possess mucolytic properties, demonstrate synergy [33], and have antimicrobial properties [35, 36] they may serve in both solubilization of mucinous secretion whilst also acting as an anti-microbial agent. Owing to BromAc^®^’s strong mucolytic activity, with its hydrolytic action on peptide and glycosidic bonds along with its disulfide reductive properties [37, 38] it is envisaged that these two agents may solubilize dense and purulent sputa secreted in the airways in disease states. Preliminary studies with BromAc^®^ on cystic fibrosis sputa has shown that it clearly disintegrates the sticky mucinous mass into a free-flowing solution *in vitro* (unpublished data) and hence in the current study we investigate the mucolytic effect of nebulised BromAc^®^ on two sputa models since we envisage using this formulation in aerosol form for treatment. The two sputa models used in this study are artificial sputa (AS) as well as simulated sputa (SS) that is mucinous secretion from PMP patients which has been specially treated to represent thick airway sputa. Nebulised delivery of therapeutics is a convenient method for treating diseases of the airways [39].

## Material and Methods

### Materials and reagents

For preparing artificial sputa the following materials were purchased from Sigma Aldrich, Sydney, Australia: Porcine mucin, Salmon sperm DNA, potassium chloride, sodium chloride, Trizma^®^ base. TPTZ [2,4,6-Tris(2-pyridyl)-s-triazine], Ferrous chloride. PMP mucin of soft grade was obtained from a clinical sample that had been assessed for its softness index [40]. Pipettes (0.5 ml), capillary tubes, endotracheal tubes size 9.0, Nebulizer equipment (Philips Respironics InnoSpire Elegance Compressor, flow rate 7 L/min, 10 psi).

### Artificial sputa preparation

Artificial sputum was prepared following protocol as detailed by Kirchner [41]. Briefly, 250 mg of porcine mucin, 200 mg of sperm DNA, 295 mg of Diethylene Triamine Penta-Acetic Acid (DTPA), 25 mg sodium chloride, 110 mg potassium chloride, 140 mg Tris base were mixed in a volume of 30 ml of distilled water and pH adjusted to 7.0 using Tris base. The volume of the solution was then adjusted to 35 ml. Further dilutions were then carried out to adjust viscosity as desired.

### Preparation of PMP Mucin as a model of sputa

Six grams of soft PMP mucin was homogenised using a shredder with phosphate buffer saline (PBS) (3.0 ml) with sonification and vortexed until a homogenous mixture was formed, with incubation at 37° C to remove air bubbles and after which viscosity was adjusted (further dilutions with PBS). The pH was adjusted to 7.0 using either 1.0 M NaOH or 0.1 N Hydrochloric acid.

### Measurement of dynamic viscosity

Dynamic viscosity (γ) of sputum was measured using the capillary tube method as outlined by Santander and Castellano [42].

### Measurement of pipette flow time

Using a 0.5 ml glass pipette fixed at an angle of 60°, 0.5 ml of sample at 25° C (ambient room temperature) was sucked up the pipette and the time taken to empty 0.3 ml of the sample was timed in sec.

Pipette flow time (€) was calculated as follows:

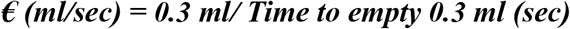

### Treatment with aerosol Acetylcysteine

#### Artificial sputa (AS)

Artificial sputa were prepared as detailed in earlier and 1.5 ml of artificial sputum was carefully emptied into an endotracheal tube (in triplicates). A volume of 6.0 ml of NAC 10 mg/ml in PBS (pH 7.0) in triplicates was aerosolised using a nebulizer and passed over the sputum samples in the endotracheal tubes kept at 37° C in a water bath Similarly the experiments were repeated using NAC 20 mg/ml in PBS (pH 7.0). The controls only received PBS (pH 7.0). The aerosol delivery time to empty 6.0 ml for each treatment was 25 min.

The dynamic viscosity of the sputa was measured before and after using the capillary tube method, whilst the pipette flow time was also measured as described earlier using a 0.5 ml glass pipette. Samples were equilibrated to 25° C before measurement.

#### PMP mucin simulated sputa (SS)

A similar investigation as above was carried out to determine a comparative dynamic viscosity and pipette flow time.

#### Treatment with nebulised Bromelain (Artificial sputa, simulated sputa)

Similar treatment as in previous experiment for NAC, was carried out using Bromelain (125 and 250 µg/ml) in PBS (pH 7.0) along with control using PBS (pH 7.0). Both dynamic viscosity and pipette flow time was measured as outlined earlier for both artificial and PMP simulated sputa models.

#### Treatment with aerosol BromAc^®^ (Bromelain + NAC) (Artificial sputa, simulated sputa)

Using a combination of NAC (20 mg/ml) with either 125 or 250 µg/ml Bromelain in PBS (pH 7.0), a similar investigation was carried out as outlined earlier and measurements were made for both artificial and simulated sputa models.

#### Measurement of Bromelain in aerosolised sputum samples

Suitable dilutions such as 1/5 and 1/10 of nebulised samples in PBS were carried out with subsequent filtration (0.44 um). To 250 µl of sample was added 250 µl of 1% azocaesin solution (prepared in distilled water). The samples were agitated at room temperature (25° C) /1 hrs, after which 1.5 ml of 1% trichloroacetic acid was added, vortexed and centrifuged at 2500 RPM. To 150 µl of supernatant in a microwell was added an equal quantity of 0.5 M Sodium hydroxide solution and the OD at 410 nm was read using a spectrometer.

A standard curve for Bromelain was generated following a similar procedure with Bromelain dilutions ranging from 200 µg/ml that was serially diluted.

#### Measurement of Acetylcysteine in sputum samples

Suitable dilution such as 1/5 and 1/10 of aerosolised samples and filtration, as above was carried out. 10 mM solutions of TPTZ were prepared in distilled water (dH_2_O).

Stock solution of NAC 10 mg/ml was prepared in PBS and pH adjusted to 7.0. Stock solution (10 mM) of Fe (III) was prepared in dH_2_O. The pH of all the reagents was adjusted to 7.0. To 125 µl of TPTZ was added 125 µl of Fe (III) solution vortex mixed and then followed by the addition of 100 µl of test solution. It was then vortexed and placed on a gentle shaker for 1 hour at ambient room temperature (25° C) until full colour (blue) developed. 2.0 ml of dH_2_O was added, vortex mixed, and the OD was measured using aliquots of 200 µl in triplicates into a 96 well plate. Blanks only contained 100 µl of dH_2_O. The OD at 593 nm was read using UV spectrometer (Shimadzu). Suitable dilutions of Acetylcysteine ranging from 200 µg/ml down was prepared for the generation of standard curve.

### Calculation of D values (%) for both the dynamic viscosity (γ) and pipette flow (€)

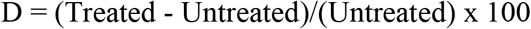

## Results

### Treatment with N-Acetylcysteine (NAC)

Treatment with PBS had very minor effect on dynamic viscosity (**γ)** of artificial sputum (AS), however, NAC at 10 and 20 mg/ml showed a marked decrease in **γ**, 6.0 and 9.8%, respectively. The pipette flow speed (**€)** indicated that treatment with PBS affected it by 20% indicating that hydration may play a substantial role on this parameter. Further, treatment with NAC 10 and 20 mg/ml also showed a corresponding increase in pipette flow (28 and 40%, respectively).

In the case of simulated sputa (SS), PBS treatment affected **γ** slightly (2%) while having almost an equal effect (16 and 17%) with treatment of NAC at the two concentration (10 and 20 mg/ml). Further there was also a corresponding increase in **€** values. Thus, both the sputa models were affected by aerosol treatment with NAC. **(Table 1. Figures 1A-D)**.

**Table 1.** shows the effect of aerosolised N-Acetylcysteine on both artificial and simulated sputa using two parametric measurements such as dynamic viscosity and pipette flow time. NAC: N-Acetylcysteine. **↓ = decrease;** ↑= increase

| DYNAMIC VISCOSITY (γ) MEASUREMENTS (cSt) |  |  |  |  |  |  |
| --- | --- | --- | --- | --- | --- | --- |
|  | ARTIFICIAL SPUTUM (AS) |  |  | SIMULATED SPUTUM (SS) |  |  |
| Treatment | Before | After | D (%) | Before | After | D (%) |
| Control | 1.382±0.013 | 1.373±0.001 | 0.65 ↓ | 1.748±0.006 | 1.713±0.004 | 2 ↓ |
| NAC 10 mg/ml |  | 1.300±0.002 | 6.0 ↓ |  | 1.467±0.011 | 16 ↓ |
| NAC 20 mg/ml |  | 1.247±0.041 | 9.8 ↓ |  | 1.453±0.055 | 17 ↓ |
| PIPETTE FLOW SPEED (€) (ml/sec) |  |  |  |  |  |  |
|  | ARTIFICIAL SPUTUM (AS) |  |  | SIMULATED SPUTUM (SS) |  |  |
| Treatment | Before | After | D (%) | Before | After | D (%) |
| Control | 0.005±0.001 | 0.006±0.001 | 20 ↑ | 0.0137±0.005 | 0.0153±0.001 | 12 ↑ |
| NAC 10 mg/ml |  | 0.0064±0.003 | 28 ↑ |  | 0.0183±0.001 | 34 ↑ |
| NAC 20 mg/ml |  | 0.007±0.001 | 40 ↑ |  | 0.02±0.002 | 46 ↑ |

**Figure 1.**
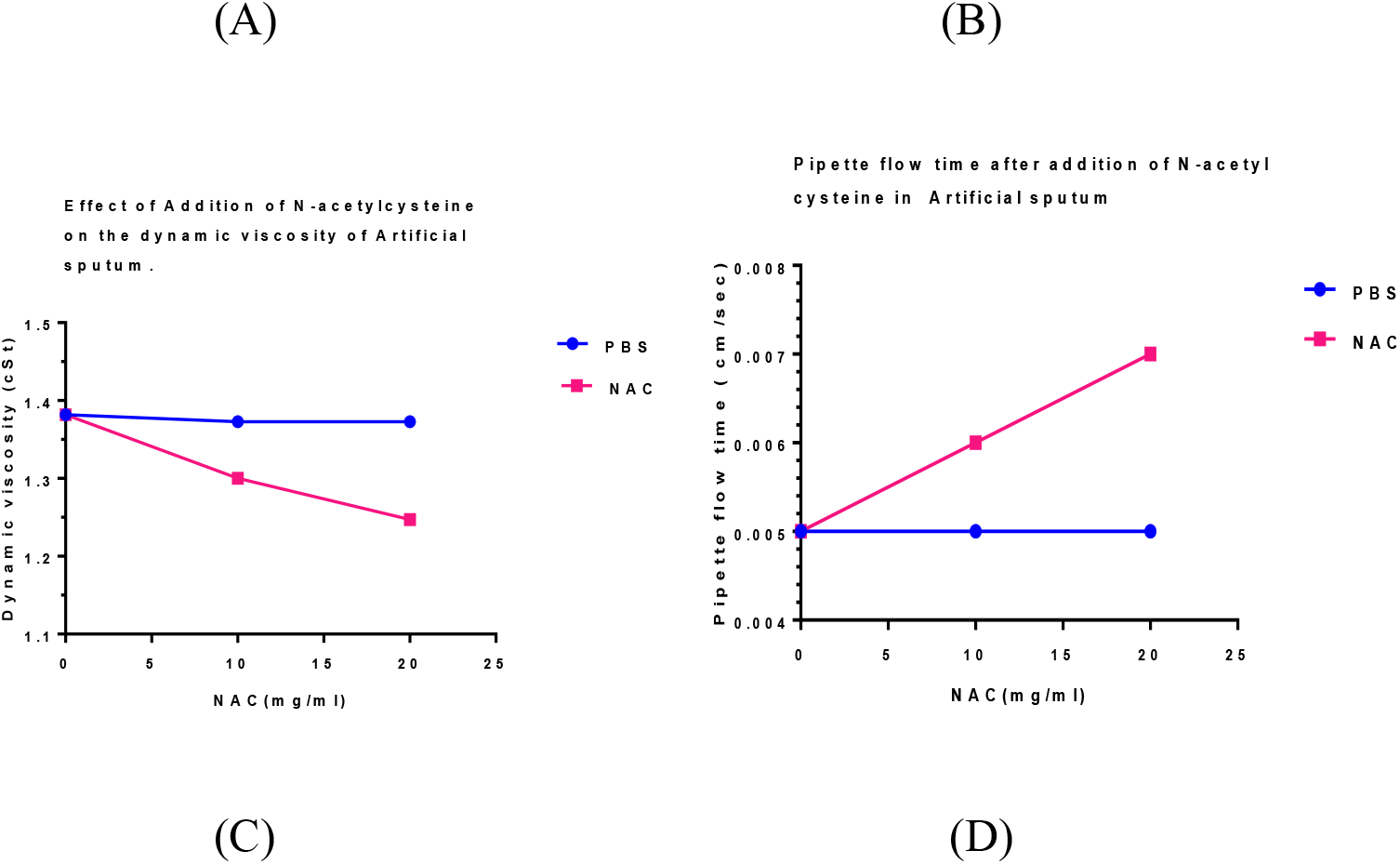
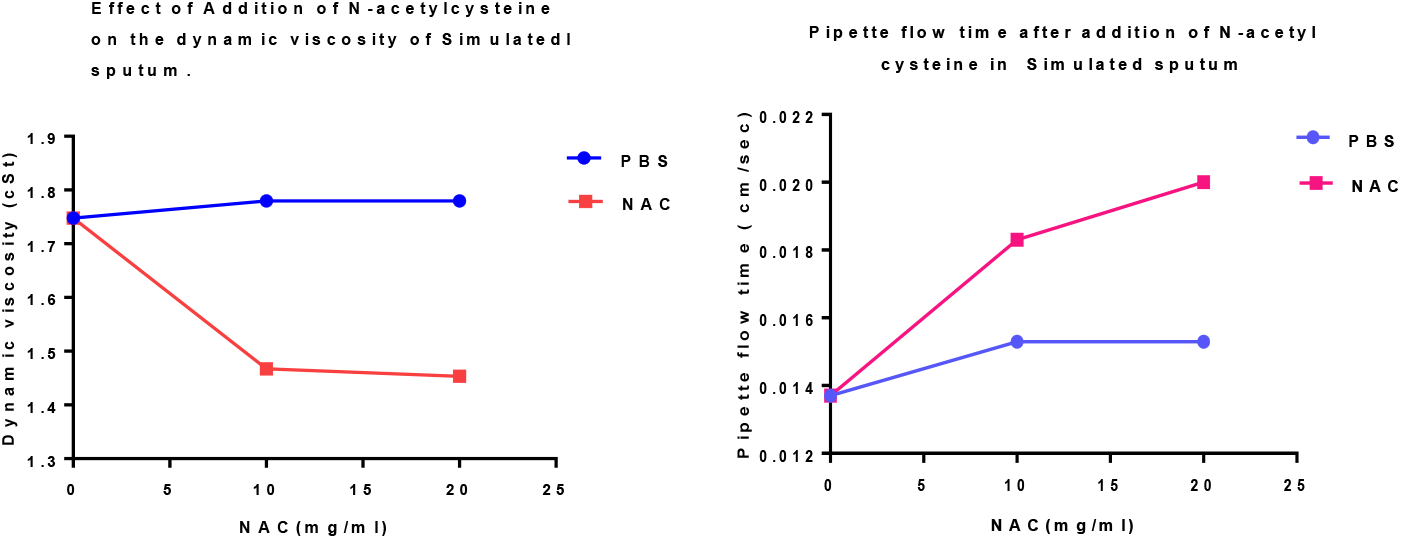
**A)** indicates that in artificial sputa (AS), with the addition of N-Acetylcysteine, there is a noticeable concentration dependent reduction of dynamic viscosity. **B)** shows that in AS there is a linear increase in pipette sputum flow with the addition of increasing amount of NAC. **C)** indicates that there is a considerable drop in dynamic viscosity of simulated sputum (SS) with the addition of NAC 10 and 20 mg/ml, as compared to control. **D)** shows the increase in pipette sputum flow after the addition of increasing amounts of NAC in SS model.

### Treatment with Bromelain (BR)

The dynamic viscosity **(γ)** of both the sputa AS and SS were slightly affected by aerosol PBS, 0.65 and 2.0% respectively, however treatment with Bromelain at 125 and 250 µg/ml indicated a noticeable drop in (**γ)** for both the sputa models, with much higher drops for AS. The effect at the two concentrations of Bromelain in SS model was almost the same. When the pipette flow speed (€) was examined, aerosol PBS showed an effect on both the sputa models 20 and 12% in AS and SS, respectively, however this effect was very high when treated with Bromelain in both the models (**Table 2 and Figures 2AD)**

**Table 2.** shows the effect of aerosolised Bromelain (BR) on both artificial (AS) and simulated sputa (SS) using two parametric measurements, dynamic viscosity (**γ)** and pipette flow time (**€). ↓ = decrease;** ↑ = increase

| DYNAMIC VISCOSITY($\gamma$) MEASUREMENTS (cSt) | | | | | | |
| --- | --- | --- | --- | --- | --- | --- |
|  | ARTIFICIAL SPUTUM (AS) |  |  | SIMULATED SPUTUM (SS) |  |  |
| *Treatment* | *Before* | *After* | *D (%)* | *Before* | *After* | *D (%)* |
| Control | 1.382±0.013 | 1.373±0.001 | 0.65 ↓ | 1.748±0.006 | 1.713±0.004 | 2 ↓ |
| BR 125 μg/ml |  | 1.012±0.022 | 27 ↓ |  | 1.612±0.004 | 8.0 ↓ |
| BR 250 μg/ml |  | 0.870±0.003 | 36 ↓ |  | 1.600±0.009 | 8.5 ↓ |
| PIPETTE FLOW SPEED (€) (ml/sec) |  |  |  |  |  |  |
|  | ARTIFICIAL SPUTUM (AS) |  |  | SIMULATED SPUTUM (SS) |  |  |
| *Treatment* | *Before* | *After* | *D (%)* | *Before* | *After* | *D (%)* |
| Control | 0.005±0.001 | 0.006±0.001 | 20 ↑ | 0.0137±0.005 | 0.0153±0.001 | 12 ↑ |
| BR 125 μg/ml |  | 0.0218±0.003 | 336 ↑ |  | 0.046±0.001 | 243 ↑ |
| BR 250 μg/ml |  | 0.0290±0.001 | 480 ↑ |  | 0.073±0.004 | 443 ↑ |

**Figure 2.**
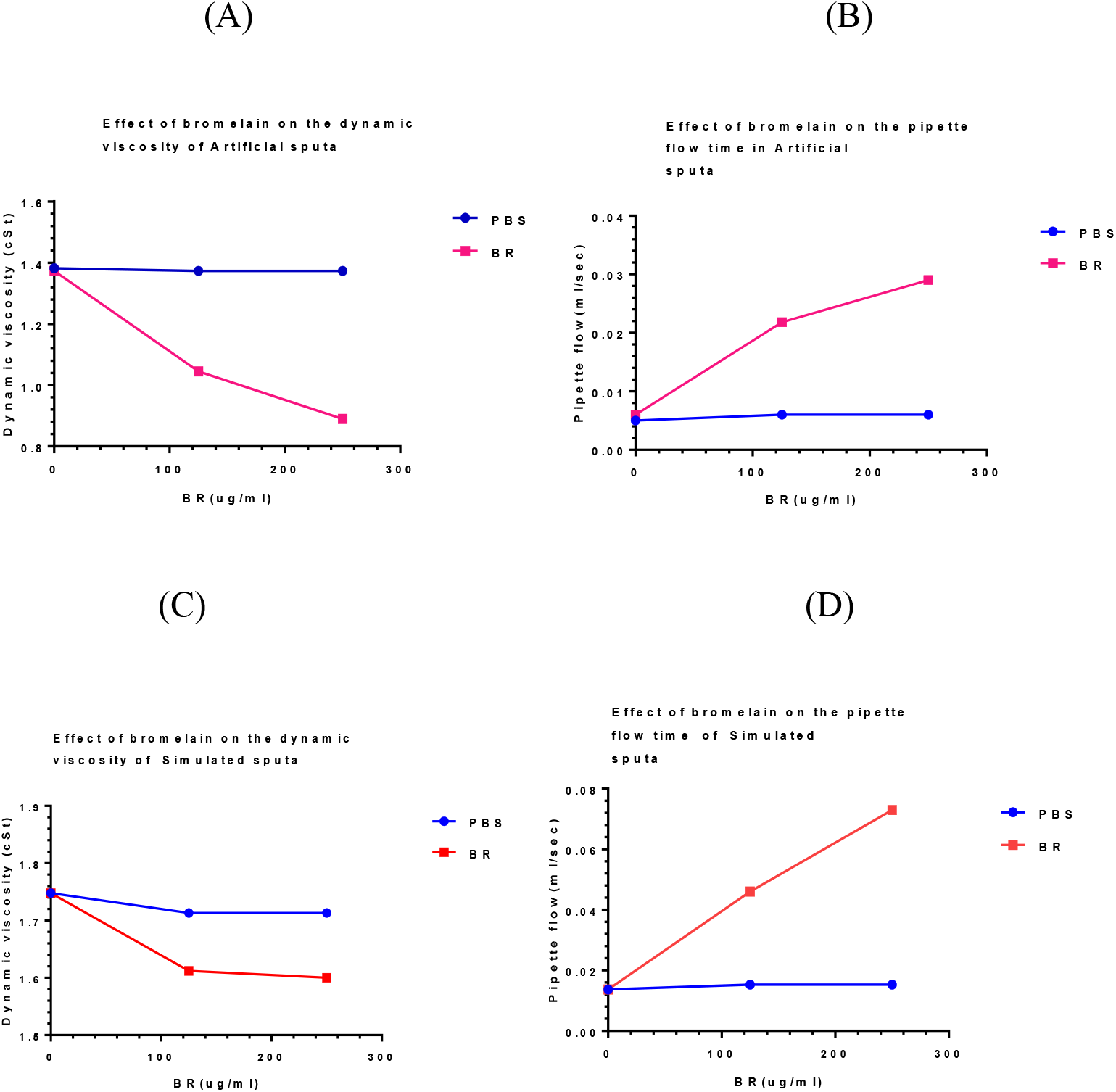
**A)** shows that in artificial sputum (AS), with the addition of Bromelain (BR) there is a considerable reduction of viscosity in comparison to control (PBS). **B)** shows that the pipette flow in AS is considerably increased as compared to control (PBS) when treated with Bromelain. **C)** shows that in simulated sputa (SS), the addition of Bromelain shows a reduction in viscosity in comparison to control (PBS). shows that in SS, the pipette flow is highly increased compared to control (PBS), after treatment with Bromelain.

### Treatment with Bromelain and Acetylcysteine (BromAc^®^)

Treatment with Bromelain at the two concentrations (125 and 250 µg/ml) with NAC 20 mg/ml had a noticeable effect on **(γ)** with sufficient decrease in both the sputa models. This was further indicated in **(€)** values with very high increase in both the models (**Table 3 and Figures 3A-D)**.

**Table 3.** shows the effect of aerosolised BromAc^®^ (Bromelain (BR) 125 or 250 µg/ml + NAC 20 mg/ml) on both artificial (AS) and simulated sputa (SS) using two parametric measurements such as dynamic viscosity (**γ)** and pipette flow time (**€). ↓ = decrease;** ↑= increase

| DYNAMIC VISCOSITY($\gamma$) MEASUREMENTS (cSt) | | | | | | |
| --- | --- | --- | --- | --- | --- | --- |
|  | ARTIFICIAL SPUTUM (AS) |  |  | SIMULATED SPUTUM (SS) |  |  |
| *Treatment* | *Before* | *After* | *D (%)* | *Before* | *After* | *D (%)* |
| Control | 1.382±0.013 | 1.373±0.001 | 0.65 ↓ | 1.748±0.006 | 1.713±0.004 | 2 ↓ |
| BR 125 µg/ml +<br>NAC 20 mg/ml |  | 0.95±0.001 | 31 ↓ |  | 1.416±0.033 | 19 ↓ |
| BR 250 µg/ml +<br>NAC 20 mg/ml |  | 0.800±0.002 | 42 ↓ |  | 1.282±0.027 | 27 ↓ |
| PIPETTE FLOW SPEED (€) (ml/sec) |  |  |  |  |  |  |
|  | ARTIFICIAL SPUTUM (AS) |  |  | SIMULATED SPUTUM (SS) |  |  |
| *Treatment* | *Before* | *After* | *D (%)* | *Before* | *After* | *D (%)* |
| Control | 0.005±0.001 | 0.006±0.001 | 20 ↑ | 0.0137±0.005 | 0.0153±0.001 | 12 ↑ |
| BR 125 µg/ml +<br>NAC 20 mg/ml |  | 0.033±0.002 | 556 ↑ |  | 0.061±0.002 | 343 ↑ |
| BR 250 µg/ml +<br>NAC 20 mg/ml |  | 0.035±0.001 | 600 ↑ |  | 0.114±0.002 | 733 ↑ |

**Figure 3.**
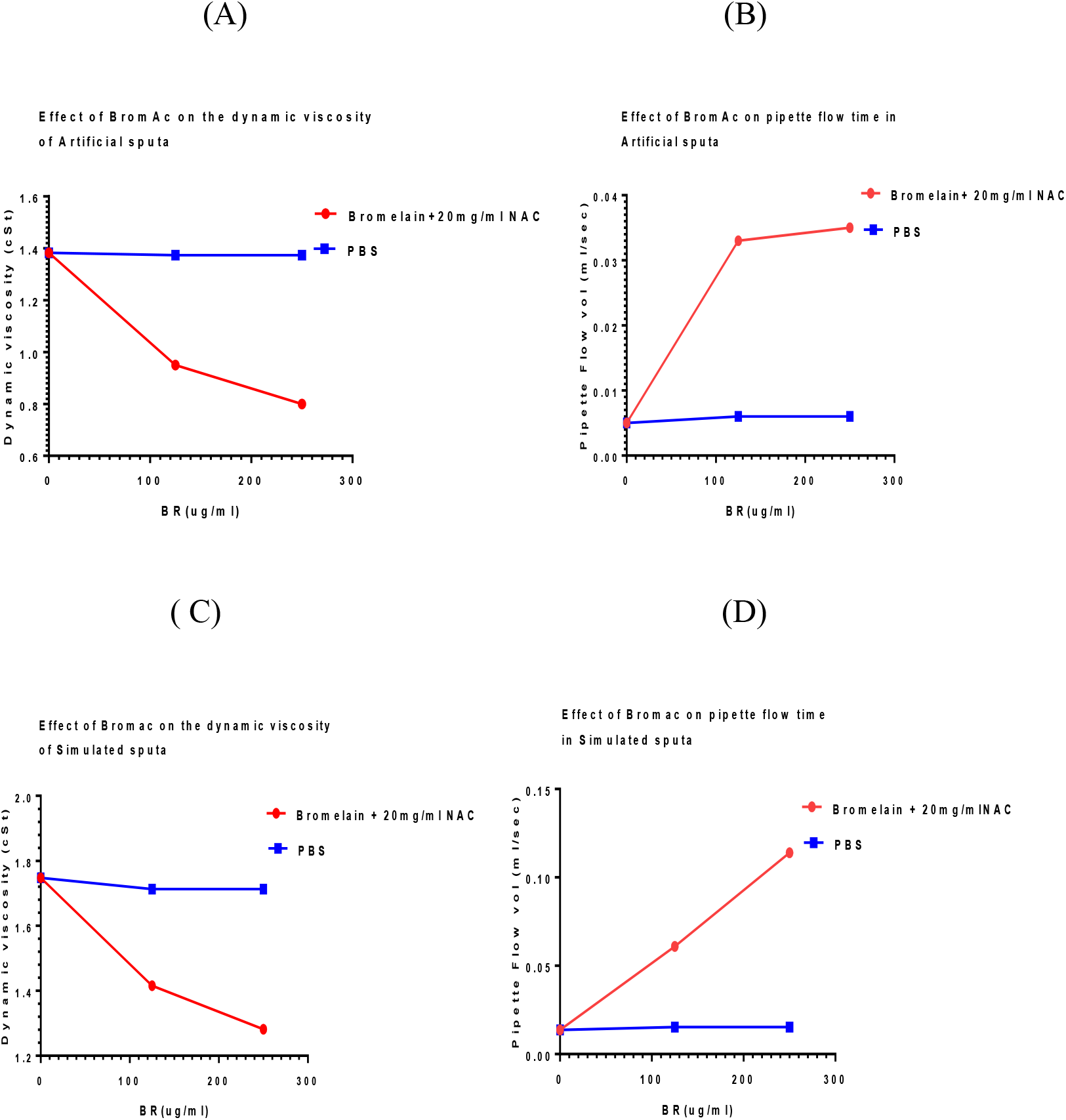
**A)** shows that in artificial sputa (AS) there is a noticeable drop in dynamic viscosity after treatment with BromAc^®^ (Bromelain 125 and 250 µg/ml + NAC 20 mg/ml). **B)** shows that in AS the flow rate increase with the addition of BromAc^®^. **C)** shows that in simulated sputa (SS) there is a considerable drop in dynamic viscosity with the addition of 125 or 250 µg/ml Bromelain + NAC 20 mg/ml. **D)** shows that in SS, there is almost a linear increase in flow rate (pipette emptying time) with the addition of increasing the amount of Bromelain and 20 mg/ml NAC.

Bromelain at 125 µg/ml as individual agent showed a concentration of 30.58 µg/ml in the AS and with 250 µg/ml it was 58.63 µg/ml indicating that doubling the concentration almost doubled the sequestered Bromelain. However, in the SS, there was very little difference between the two Bromelain concentrations (57.91 vs 61.64 µg/ml). NAC as individual agent showed only a small difference in concentration between 10 and 20 mg/ml in AS whilst slightly larger difference in SS models. When Bromelain was delivered in NAC 20 mg/ml, Bromelain concentration in the 125 µg/ml was half of that found in the 250 µg/ml Bromelain in the AS model whilst in the SS model, there was only a 11% difference. Although NAC 20 mg/ml was delivered with either 125 or 250 µg/ml Bromelain, NAC analysis indicated that in the AS model, the difference was small with a slightly larger difference in the SS model **(Table 4, Figures 4A-C)**.

**Table 4.** shows the concentration of either Bromelain (BR) or Acetylcysteine (NAC) in two sputa models (AS and SS) after passage of nebulised solution over the sputa with either just Bromelain, NAC, or their combination over 25 min.

|  | *ARTIFICIAL SPUTA (AS)* |  | *SIMULATED SPUTA (SS)* |  |
| --- | --- | --- | --- | --- |
| **Reagent** | **BR (µg/ml)** | **NAC (mg/ml)** | **BR (µg/ml)** | **NAC (mg/ml)** |
| BR 125 µg/ml | 30.58±0.211 |  | 57.91±0.562 |  |
| BR 250 µg/ml | 58.63±0.89 |  | 61.64±0.88 |  |
| NAC 10 mg/ml |  | 2.75±0.143 |  | 1.85±0.310 |
| NAC 20 mg/ml |  | 2.90±0.210 |  | 2.367±0.133 |
| BR 125 µg/ml +<br>NAC 20 mg/ml | 41.23±1.212 | 2.41±0.122 | 77.56±3.12 | 1.62±0.093 |
| BR 250 µg/ml +<br>NAC 20 mg/ml | 79.15±2.22 | 2.22±0.184 | 88.21±4.11 | 2.16±0.132 |

**Figure 4.**
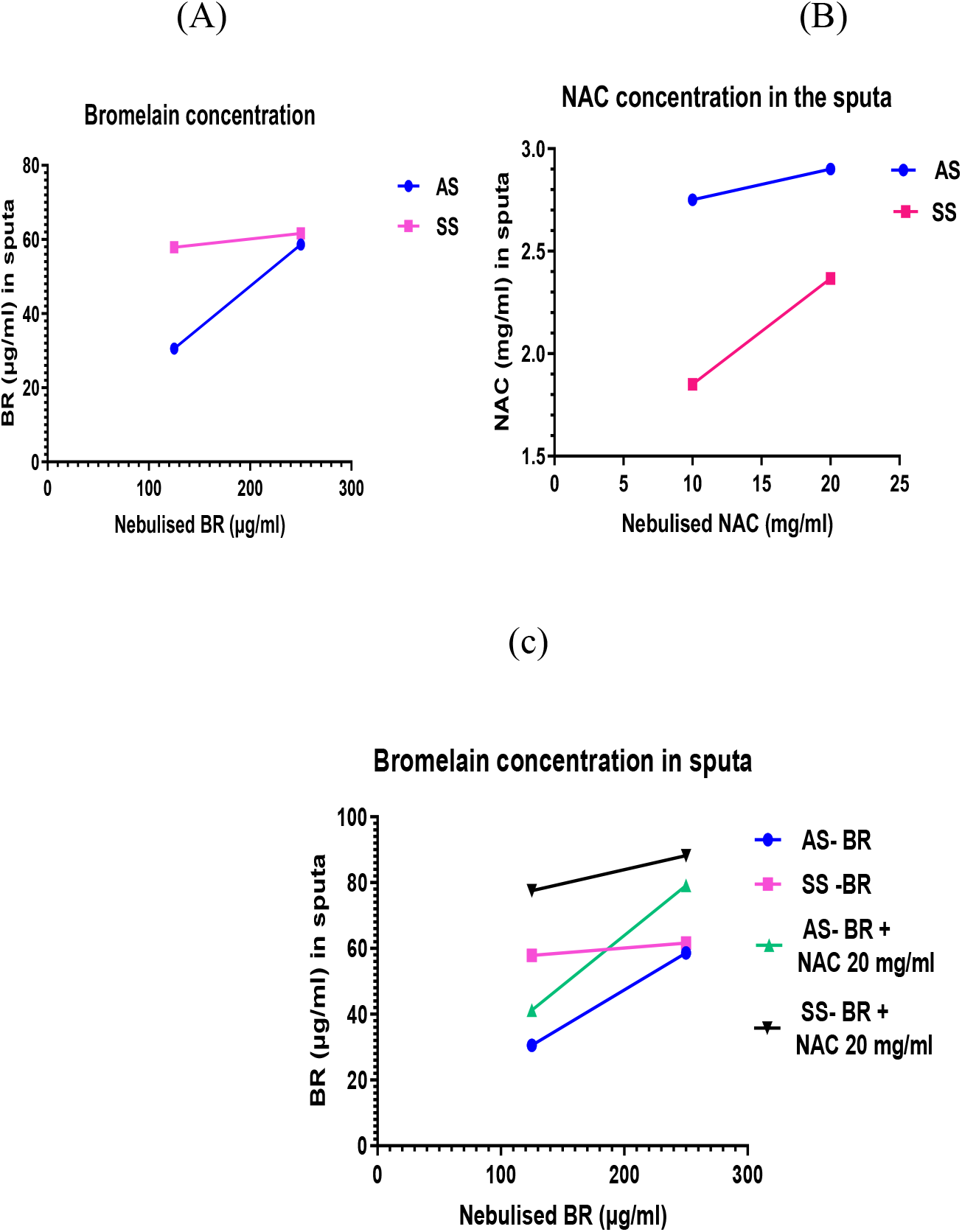
**A)** shows that the concentration of Bromelain in the artificial sputa (AS) increases almost 2-fold with exposure to two-fold increase in concentration of aerosolised Bromelain (125 vs 250 µg/ml). However, the difference was relatively small for the SS, simulated sputa 58 vs 62 µg/ml. **B)** shows that the difference between the low (10 mg/ml) and the high (20 mg/ml) NAC in the artificial sputa (AS) was relatively small (difference of 0.15 mg/ml) when exposed to aerosolised NAC. However, in the simulated sputa (SS), the difference was relatively larger (1.85 vs 2.37 mg/ml). about 28% increase. **C)** shows comparative levels of Bromelain in artificial and simulated sputa before and after addition of 20 mg/ml NAC to the aerosolised solution. The concentration of Bromelain is relatively higher in both the sputa models, in the presence of NAC 20 mg/ml.

Further, we compared both the D values of both dynamic viscosity (γ) and flow speed (€) between artificial and simulated sputa (Table 5 and Figure 5). It has been shown that small changes in viscosity of the mucinous solution can produce a large increase in flow rate based on pipette emptying time. This may have a bearing on ciliary clearance that needs further investigation.

**Table 5.** Comparison of Dynamic viscosity (γ) and Flow speed (€) between the two sputa models when treated with Bromelain 125 or 250 µg/ml and in combination with NAC 20 mg/ml. shows a comparison of D values (for both the dynamic viscosity (**γ)** and flow speed (**€**) between artificial sputa (AA) and simulated sputa (SS) treated with Bromelain only and with the addition of NAC 20 mg/ml. With the addition of NAC 20 mg/ml, there is a slight increase in (**γ)** in AS whilst the difference was much larger in SS model. Further, a similar trend in (€) was seen in both the sputa.

| **D value (%)** |  |  |  |  |
| --- | --- | --- | --- | --- |
|  | *Artificial Sputa (AS)* |  | *Simulated Sputa (SS)* |  |
| **Treatment** | **Dynamic Viscosity ($\gamma$)** | **Flow Speed ($\epsilon$)** | **Dynamic Viscosity ($\gamma$)** | **Flow Speed ($\epsilon$)** |
| 125 μg/ml BR | 27 | 336 | 8.0 | 243 |
| 250 μg/ml BR | 36 | 480 | 8.2 | 443 |
| 125 μg/ml BR + NAC 20 mg/ml | 31 | 556 | 19 | 343 |
| 250 μg/ml BR + NAC 20 mg/ml | 42 | 600 | 27 | 733 |

**Figure 5.**
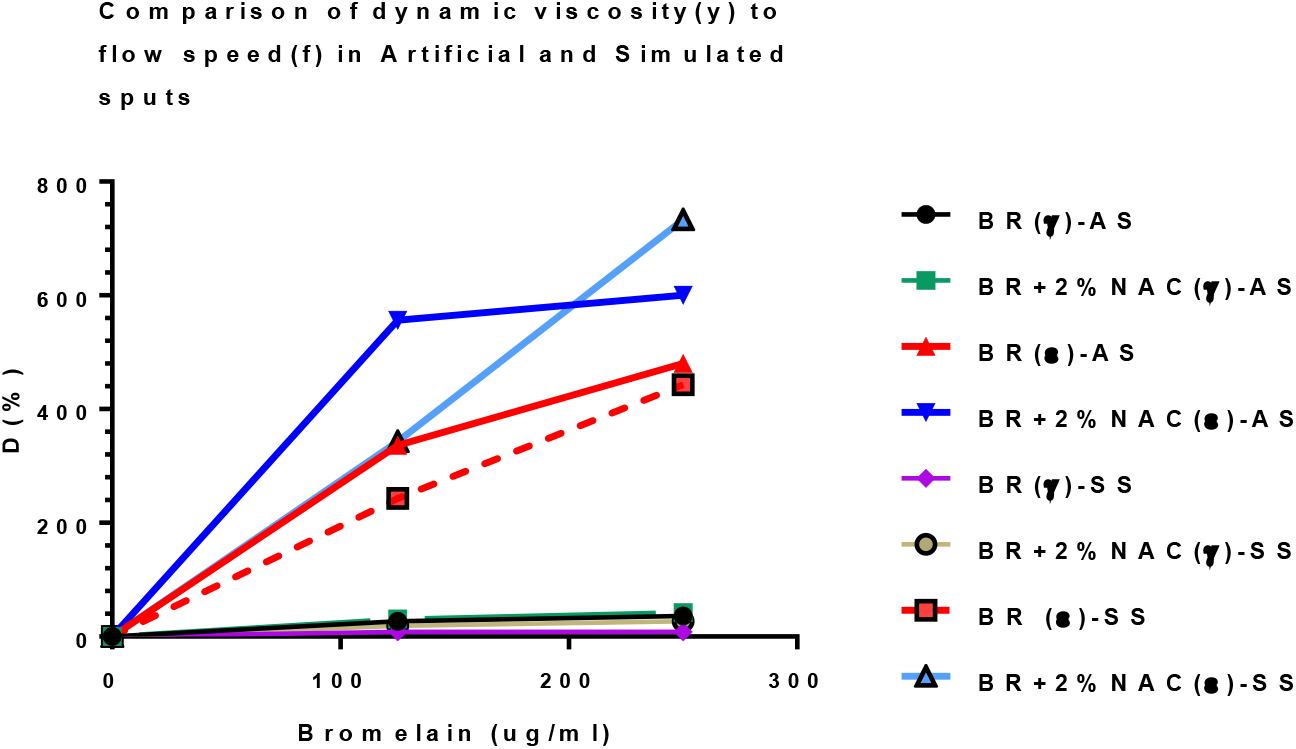
shows the relative differences in enhancement D (%) of the two parameters measured such as dynamic viscosity (**γ**) and flow speed **€** in the artificial sputum (AS) and simulated sputum (SS), when treated with either Bromelain (BR) 125 or 250 µg/ml alone or in combination with NAC 20 mg/ml (BromAc^®^). Viscosity changes in the different groups are amplified in flow speed showing the effect of viscosity on the latter. Treatment with BromAc has a much higher effect on flow speed compared to treatment with bromelain alone. D = (Treated - Untreated)/(Untreated) x 100.

Although, with the addition of NAC 20 mg/ml to Bromelain 125 and 250 µg/ml in artificial sputum, the decrease in viscosity was only 4-6 % compared to Bromelain alone (from 31-42% to 27-36%, respectively), there was a very large increase in flow speed (€) indicating that a small decrease (changes) in viscosity may affect clearance of the fluid. The high BromAc concentration group indicates almost twice reduction of pipette emptying time compared to the low BromAc group. This does not correlate with their respective dynamic viscosity (γ) that may indicate that highly glycosylated mucin may affect the viscosity to a significant level. In simulated sputa, with the addition of NAC 20 mg/ml to Bromelain 125 and 250 µg/ml, the decrease in dynamic viscosity was 19 and 27% compared to 8 and 8.2%, respectively. Further, a similar trend in flow speed (€) was seen in both sputa.

## Discussion

Since, clearance of airway secretion, is mainly dependent on its rheological parameters, we decided to measure the dynamic viscosity (**γ)** and the flow speed (€) of the sputa before and after treatment with BromAc^®^ using two model sputa: artificial sputa (AS) and simulated sputa (SS), specially formulated to represent thick and static sputa. However, since two different agents (Bromelain and Acetylcysteine) are incorporated into the formulation (BromAc^®^), the effect of individual agents were first investigated with subsequent studies on their combination. The differences for both **γ** and € (pre-treated as opposed to treated) were calculated as a percentage denoted by D. Additionally, we also investigated the sequestration of Bromelain and NAC in the sputa before and after aerosol delivery for individual agents as well as their combinations since we wanted to correlate the concentrations of the agents within the sputa with the changes in rheological properties. Treatment of artificial sputa (AS) with aerosolised PBS indicated a minute drop in **γ** (0.65%) that may be mainly due to hydration, whilst in both the NAC 10 and 20 mg/ml, the reduction was by 6.0 and 10%, respectively. NAC is a well-known antioxidant and hence the reduction of disulfide linkages found in the protein and mucin components may be responsible for this change in viscosity [43, 44]. In comparison, simulated sputa (SS) in PBS showed a 2.0% drop in **γ** with substantial effect in NAC 10 and 20 mg/ml (**γ** =16 and 17%, respectively). This difference between both the sputa may be due to their differences in composition and variability of the constituents. The SS is mainly composed of mucinous mass that is heavily glycosylated with cellular debris and other components that may contain abundant disulfide linkages [45] and hence highly prone to the reductive action of NAC. Since the action of NAC is substantial in simulated sputa (SS) it may indicate that the percentage of disulfide linkages within the matrix may be much higher compared to artificial sputa (AS) (at NAC 20 mg/ml, γ was 17 and 10%, respectively).

The effect on flow speed **€** was substantial (20% increase) with PBS treatment, indicating that hydration alone may have considerable effect on the viscoelastic property of sputum in AS which is agreement with other researchers on rheology of sputa [5, 46]. Further, at NAC 10 and 20 mg/ml, the effect was 28 and 40% respectively. In, comparison, the SS displayed a substantial increase in flow speed **€** with only PBS (12%) whilst both the NAC 10 and 20 mg/ml had a much higher impact (34 and 46% increase). The differences between the two sputa models in flow speed may be mainly due to their differences in composition. Most likely, the percentage of disulfide bonds in the SS model may be much higher compared to AS model and hence their reduction by NAC shows a considerable effect on this parameter. Further studies to determine the concentration of disulfide bonds in both the sputa before and after treatment may enable an understanding of the effect and the degree to which these bonds are affected and hence the observed effect on rheological parameters.

Treatment with Bromelain showed a marked drop in **γ** in both the treatment groups (0.125 and 250 µg/ml) with a drop of 27 and 36%, respectively in AS model with substantial effect on **€** (336 and 480%, respectively) indicating the impact of dynamic viscosity on the flow speed of sputa and ciliary clearance [47]. In the case of SS model again there was drop in **γ** of 8.0 and 8.2% for the two concentrations of Bromelain (125 and 250 µg/ml), with almost no difference. This similarity in **γ** may be due to the high glycosylation and high mucin present with the sputa. However, the effect of Bromelain on γ amplified the effect on the **€** values as indicated (243 and 443%). Hence, Bromelain seems to affect both the parameters monitored showing that its hydrolytic properties on proteins and glycoproteins affect the rheological properties of both the sputa models. The variation in efficacy between the two models may be attributed to their differences in composition. In comparison to NAC, Bromelain has a much greater effect on the two rheological parameters monitored in AS, whilst the impact although less on **γ**, had much higher impact on € values for the SS model, indicating that the enzymic reactions on these models’ sputa shows a much higher activity with greater depolymerisation effect that affected the parameters monitored. Of note, in the SS model, the effect on **γ** values were higher in the NAC treated compared to Bromelain which may indicate the high disulfide content in the sample sputa. This needs further verification; however, studies have shown the depolymerising effect of NAC through the reduction of these vital linkages that dictates the molecular geometry of proteins [44, 48].

When the sputa were treated with BromAc^®^ (Bromelain 125 and 250 µg/ml + NAC 20 mg/ml), γ was considerably affected as shown by the percentage differences between untreated and treated sputa in AS for both the 125 and 250 µg/ml Bromelain with values of 36 and 42%. Further this effect was amplified in the € parameter with corresponding 556 and 600%. In the case of SS, the **γ** parameter between 125 and 250 µg/ml Bromelain showed 19 and 27%, with a substantial difference in € values, (343 and 733%, respectively). Hence, the effect on dynamic viscosity was considerably amplified in € with considerable rise because of treatment with Bromelain plus NAC 20 mg/ml. These variations between the two models may be mainly due to their variability in composition that needs further studies.

The concentration of Bromelain in the sputa after treatment indicated that with Bromelain alone, AS showed double the concentration when treated with 250 µg/ml compared to 125 µg/ml Bromelain (58.63 vs 30.58 µg/ml, respectively) with some correlation to their observed effect on **γ**. On the contrary, the concentration of Bromelain in SS was almost similar in both the Bromelain groups (57.91 vs. 61.64 µg/ml), with correlation with their activity, further showing that Bromelain may have accumulated to perhaps saturation in the models over 25 min of aerosol delivery. The differences between the two sputa may also be related to their heterogenous composition. In the case of NAC, the concentration measured in the AS sputa seems to correlate with their activity as measured by dynamic viscosity, and like wise for the SS model. A similar correlation was seen with €.

Finally, the analysis for Bromelain in the BromAc^®^ (Bromelain + NAC 20 mg/ml) indicated that there was a corresponding double the concentration in the 250 µg/ml Bromelain as opposed to 125 µg/ml Bromelain (79 vs 41 µg/ml Bromelain) in the AS model with correlation to their activity. This was not the case with SS model that showed only about 11 µg/ml less in the 125 µg/ml group (78 vs 88 µg/ml) indicating that near saturation of Bromelain may have taken place. Finally, in the BromAc^®^ group, the NAC sequestered was almost similar in both the high and low Bromelain group since NAC 20 mg/ml was delivered in both the groups in the AS model. However, there was a difference in the SS model with the 125 µg/ml Bromelain having less NAC concentration (1.62 vs 2.16 mg/ml). Hence, these fluctuations may be partly due to differences in composition.

Dynamic viscosity **(γ)** and flow speed (€) on a comparable basis indicates that that both the sputa models were affected. The **γ** in SS was affected less by Bromelain as compared to AS and this may be due to their differences in composition, the former having a higher protein and glycosidic linkages compared to the latter. However, when treated in combination with NAC 20 mg/ml as compared to Bromelain alone, the effect on γ was higher in SS indicating that NAC may be playing a crucial role in reduction of disulfide bridges in the sample affecting the rheology of the sputum. On the other hand, when examining, €, both the sputa were well affected. There is a marked difference in € between Bromelain and that in combination with NAC indicating the importance of NAC in depolymerising the sputa. Importantly, these results further emphasise the high impact of these agents (Bromelain, NAC, BromAc^®^) on γ and € such that any slight increase of the former magnifies the latter. This finding is important particularly in developing formulas for improving the flow and clearance of sputum from the lungs.

Although the effect of the individual agents and their combinations showed clearly that that they have substantial effect on both the sputa models with variable effects on the two parameters measured, the differences between the models observed may be mainly may be due to their heterogenous composition. The Artificial sputa was laboratory formulated containing mainly porcine mucin, DNA, and a number of other additives, that did not include hyaluronic acid found in cystic fibrosis and COVID-19 sputa. On the other hand, the simulated sputa (SS) contained a heavy load of mucinous materials which are substantially glycosylated (-O- linkages) that are prone to the enzymic action of Bromelain, whilst having a high load of cellular materials, lipids, sialic acid etc. in addition [45]. However, the observed rheological effects in these two model sputa may be sufficient to indicate that aerosol delivery of BromAc^®^ will have considerable effect on patient sputum from CF, COVID-19 or other sputum producing respiratory disease. Recent studies have indicated that Cystic fibrosis and COVID-19 airway secretion are heavily loaded with double stranded DNA (>600 µg/ml) and hyaluronic acid (7.0 µg/ml) [18]. In comparison the artificial sputa (AS) used in the present study contained about 5714 µg/ml DNA and 7142 µg/ml mucin. Hence, the variability in COVID-19 and CF sputum should not present any barrier to BromAc^®^ since Bromelain will hydrolyse the β-1-4 glycosidic linkages found in hyaluronic acid since it has been widely used in hydrolysis of chitosan, a complex heavily glycosylated glycoprotein [37]. In addition, earlier studies using cystic fibrosis sputa showed that BromAc^®^ reduced the thick sputum within the hour to a free-flowing liquid that used BromAc^®^ in solution instead of aerosol (unpublished data). Importantly, CF and COVID-19 sputa share very common chemical identity since they have similar composition [18]. Further, the SS model used in this study is heavily glycosylated with abundant glycosidic bonds and the mucolytic efficacy of BromAc^®^ in these sputa is a clear indication of its efficacy on chemical compounds containing the β1-4 glycosidic linkages that is found in hyaluronic acid. We and others see the potential of mucus targeting in COVID-19 [49]. Although the current study is very promising in the development of BromAc^®^ for application in CF and COVID-19, further investigation using patient sputum with aerosol BromAc^®^ is warranted.

## DATA AVAILABILITY

The datasets generated during and/or analysed during the current study are available from the corresponding author on reasonable request.

